# Evolutionary impact of codon specific translation errors at the proteome scale

**DOI:** 10.1101/2022.10.11.511697

**Authors:** Cedric Landerer, Jonas Pöhls, Agnes Toth-Petroczy

## Abstract

Errors in protein synthesis can lead to non-genetic phenotypic mutations, which contribute to generating a wide range of protein diversity. There are currently no methods to measure proteome-wide amino acid misincorporations in a high-throughput fashion, limiting their detection to specific sites and few codon-anticodon pairs. Therefore, it has been technically challenging to estimate the evolutionary impact of translation errors. Here, we developed a computational pipeline, integrated with a novel mechanistic model of translation errors, which can detect translation errors across organisms and conditions. We revealed hundreds of thousands of amino acid misincorporations and a rugged error landscape in datasets of *E. coli* and *S. cerevisiae*. We provide proteome-wide evidence of how codon choice can locally reduce translation errors. Our analysis indicates that the translation machinery prevents strongly deleterious misincorporations while allowing for advantageous ones, and the presence of missing tRNAs would increase codon-anticodon cross-reactivity and misincorporation error rates.

## Introduction

Genetic information is processed with a high fidelity that is essential for cellular life, yet it is not exact. Despite the importance of proteins, their production is an error-prone process with error rates far exceeding genetic mutation rates. Mutations that are due to errors in transcription and translation are collectively termed phenotypic mutations (Bratulic et al.,2017; Bürger et al., 2006; Goldsmith and Tawfik, 2009; Yanagida et al., 2015). Transcriptome-wide transcription error estimates range from ∼10^−6^ in *C. elegans* (Gout et al., 2013), *E. coli, B. subtilis*, and *A. tumefaciens* (Li and Lynch, 2020) to ∼10^−5^ in *S. cerevisiae* (Gout et al., 2017; Reid-Bayliss and Loeb, 2017), and *M. florum* (Li and Lynch, 2020). Translation error rates are often measured for individual constructs (Drummond and Wilke, 2009; Gromadski and Rodnina, 2004; Kramer and Farabaugh, 2007; Loftfield, 1963; Loftfield and Vanderjagt, 1972; Mordret et al., 2019). Only one study measured translation error rates proteome-wide and found stark differences between codons in *E. coli*, ranging from ∼10^−4^ to ∼10^−3^ amino acid misincorporation per codon (Mordret et al., 2019). These error rates imply that more than 15% of all proteins contain at least one, potentially harmful, wrong amino acid (Drummond and Wilke, 2009). Translation error could thus lead to a large variation in the population of any type of protein, with most proteins having the canonical sequence and some containing random amino acid misincorporations. Since most genetic mutations are destabilizing (Eyre-Walker and Keightley, 1999; Eyre-Walker et al., 2002; Makałowski and Boguski, 1998), phenotypic mutations might also have a deleterious effect on individual protein functionality.

Previous studies have shown that organisms can employ different strategies to minimize the impact of errors during protein production: they may either regulate protein production or accumulate stabilizing mutations (Bratulic et al., 2015; Goldsmith and Tawfik, 2009) to buffer the potential deleterious effects of phenotypic mutations. While previous experiments illustrated the impact of lower than endogenous transcription and translation fidelity on a specific gene, it is still unknown if organisms face selection to improve overall translation fidelity as a global solution via costly increase of ribosome proofreading, or if translation errors are mitigated locally (Kurland, 1992).

Translation fidelity is ensured via structural and kinetic interactions between the ribosome, its co-factors, and the tRNA (Blanchard et al., 2004; Budkevich et al., 2011). These interactions serve as proofreading mechanisms giving the ribosome the ability to avoid amino acid misincorporations by rejecting non-synonymous tRNAs. However, if the correct tRNA does not arrive at the ribosome in a timely manner and no tRNA is incorporated, the ribosome will stall due to an empty A-site (Schuller and Green, 2018). Stalling can lead to frameshifts or be prevented by incorporating a non-synonymous tRNA, which results in an amino acid misincorporation (Schuller and Green, 2018). Amino acid misincorporations are rare events randomly distributed across the proteome introducing a large variety to a given protein’s population. It is currently unclear how such variety will affect the selection on a protein and how this variation is mapped onto the fitness landscape.

The evolutionary impact of amino acid misincorporations is dependent on their frequency. Experimental measurements provide data on select genes only or on a snapshot of proteomewide amino acid misincorporation. In contrast, theoretical models can provide error estimates for all sites. Previous studies calculated error rates from general kinetic ribosome parameters and tRNA abundances (Fluitt et al., 2007; Shah and Gilchrist, 2010). These models distinguish between tRNAs solely based on their abundance and their classification as (near-) cognate or non-cognate, therefore, they cannot provide any insights into specific ribosome/tRNA interactions.

Here, we developed a mechanistic model of amino acid misincorporation (Figure 1a) based on tRNA competition fitted to proteome wide amino acid misincorporation data, to study the <u>m</u>ultinomial <u>T</u>ranslation <u>E</u>rror <u>L</u>andscape (mTEL) and quantify the evolutionary impact of translation errors. We focus specifically on the interactions between tRNAs and codons in order to capture the essential codon/anticodon binding. mTEL incorporates tRNA abundances from RNAseq data and tRNA binding affinities that we derive from experimentally detected amino acid misincorporations. We developed a high-throughput pipeline (Figure 1c) to reveal the <u>e</u>mpirical <u>T</u>ranslation <u>E</u>rror <u>L</u>andscape (eTEL) by identifying amino acid misincorporations in mass-spectrometry datasets, inspired by Mordret et al., 2019. We identified a large number (up to ∼3500 in a single *S. cerevisiae* dataset) of amino acid misincorporations in 48% of examined *S. cerevisiae* datasets and 55% of examined *E. coli* datasets in the PRIDE repository (Jones and Côté, 2008; Perez-Riverol et al., 2018, 2022).

**Figure 1:**
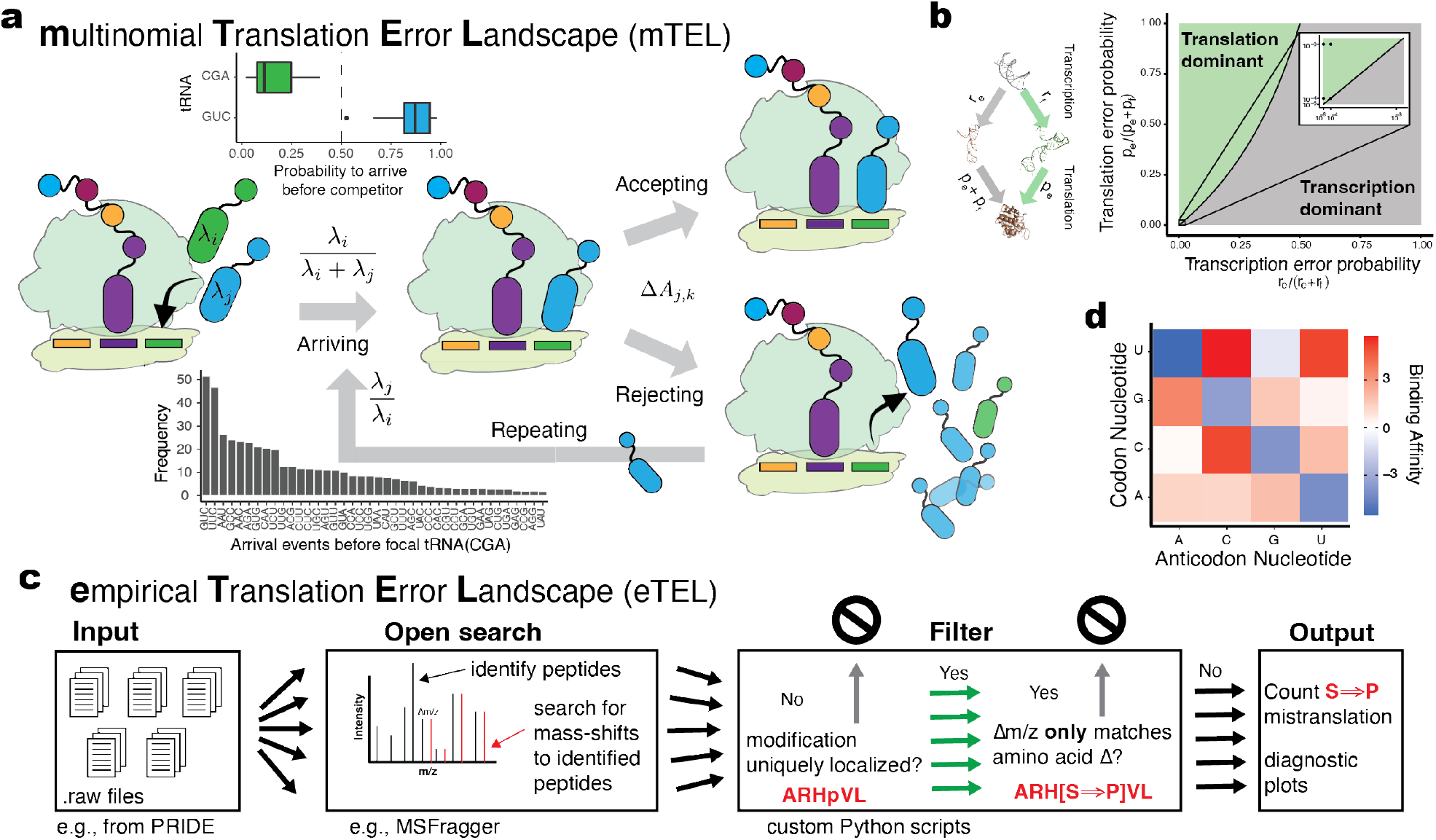
Modeling translation errors based on observed amino acid misincorporations in mass spectrometry data. **a)** The <u>m</u>ultinomial <u>T</u>ranslation <u>E</u>rror <u>L</u>andscape (mTEL) model is a mechanistic model of tRNA misincorporations. The probability of tRNA acceptance is controlled by competition for arrival at the ribosome determined by tRNA abundance and the binding affinity between codon and anticodon. Differential arrival of tRNAs is modeled as an exponential competition to arrive before another tRNA. Differences in the probability to arrive before a competitor can be substantial (top inset): the focal Ser-tRNA^CGA^ is unlikely to arrive before the competing Asp-tRNA^GUC^. Binding affinities between codon and anticodons determine the acceptance/rejection probability and are estimated from observed amino acid misincorporations. The bottom inset shows a bar graph how frequently each tRNA is expected to arrive before the focal Ser-tRNA^CGA^. **b)** Error probability space resulting from error prone protein production indicating when transcription or translation errors are more likely. Known literature estimates (Gout et al., 2017; Mordret et al., 2019; Shaw et al., 2002) suggest that observed amino acid misincorporations are most likely caused by translation errors. **c)** High-throughput pipeline for the detection of the <u>e</u>mpirical <u>T</u>ranslation <u>E</u>rror <u>L</u>andscape (eTEL) based on existing mass-spectrometry datasets (see also Figure S1). **d)** Posterior mean of binding affinities of individual nucleotides in the codon and anticodon. Watson-Crick codon/anticodon pairs show high binding affinity (blue) (see also Figure S2).

When calculating the expected fitness effects of amino acid misincorporations, we observed that their severity decreases with error probability, indicating selection against deleterious misincorporations. Combining the error frequencies and the fitness effects of amino acid misincorporations in individual proteins revealed that at the large majority of sites, amino acid misincorporations are neutral in both *S. cerevisiae* and *E. coli*. Therefore, it is likely that neither *S. cerevisiae* nor *E. coli* faces strong selection pressure to further increase translation fidelity.

In summary, we present the first data-driven, proteome-wide assessment of translation errors. We combined eTEL and mTEL into a general workflow for the <u>de</u>scription of the <u>T</u>ranslation <u>E</u>rror <u>L</u>andscape (deTEL) that is widely applicable to other organisms and conditions.

## Results

### Mechanistic modeling of tRNA incorporation allows exploration of the translation error landscape

We present a probabilistic model of tRNA incorporation as a function of tRNA arrival rates and codon/anticodon binding affinity (Figure 1a, see Methods). In short, we consider tRNA incorporation as a multi-step process: First, a tRNA has to arrive at the ribosome. Here we consider the probability of two cases: that a given tRNA arrives first, or that other tRNAs arrive beforehand and have to be rejected. We assume that tRNAs move through a cell by diffusion only, making expected arrival times of tRNAs at the ribosome proportional to their abundance (Fluitt et al., 2007; Weinberg et al., 2016). Assuming exponentially distributed arrival times, the probability that tRNA *i* will arrive before tRNA *j* is given by their expected arrival times λ:

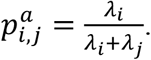

This relationship can be extended to all potential subsets of tRNAs, e.g., synonymous and non-synonymous tRNAs by defining λ_*i*_ and λ_*j*_ as the sum of the arrival rates over the respective sets.

The probability of binding between codon *i* and anticodon *j* is proportional to their affinity 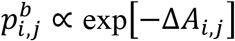. The binding affinity Δ*A*_*i,j*_ is calculated as the sum over the individual nucleotide interaction, weighted by position (see Methods for details). Describing codon/anticodon affinities as a linear combination of nucleotide affinities and position specific parameters has two main advantages. We can reduce the number of parameters and leverage information about nucleotide interaction across multiple sites and vice versa. Additionally, we can infer tRNA misincorporations without any observations of the corresponding amino acid misincorporation and provide a general model.

### Translation errors dominate transcription errors

Mass-spectrometry alone is unable to distinguish if amino acid misincorporations are caused by transcription or translation errors. While transcription errors are estimated to be orders of magnitude lower than translation errors, transcription errors are further amplified by translation. It is unclear what proportion of the measured amino acid misincorporations originate from transcription. Therefore, we explored the likelihood of the detected amino acid misincorporations to be caused by translation errors or transcription errors (Figure 1b). We modeled protein synthesis via error prone production and explored combinations of literature estimates of transcription and translation error rates leading to amino acid substitutions (see Methods). While the amplification of transcription errors will eventually lead to a transcription-error dominated system, this is limited to high transcription error rates (Figure 1b). Only if we assume the lowest translation misincorporation rate combined with the highest transcription misincorporation rate, we observe a transcription dominated system. Therefore, we assume that most amino acid misincorporations observed via mass spectrometry result from translation rather than transcription errors.

### Proteome-wide amino acid misincorporations detected in hundreds of massspectrometry datasets

While amino acid misincorporation have been explored individually for decades via reporter constructs (Drummond and Wilke, 2009; Gromadski and Rodnina, 2004; Kramer and Farabaugh, 2007), proteome wide amino acid misincorporation identification is only achievable with mass spectrometry. However, mass spectrometry data sets have large variation in the identified peptides and their frequencies, due to differences in experimental and technical conditions. In order to capture this variation, it is important to consider a large number of mass-spectrometry datasets. We applied our new high-throughput pipeline to reveal the <u>e</u>mpirical <u>T</u>ranslation <u>E</u>rror <u>L</u>andscape (eTEL) on a large number of mass-spectrometry datasets for *S. cerevisiae* downloaded from the PRIDE repository (Jones and Côté, 2008; Perez-Riverol et al., 2018, 2022). Our high-throughput pipeline, eTEL is based on MSFragger’s (Kong et al., 2017) open search algorithm (Yu et al., 2020) and uses stringent quality filters to avoid the misidentification of mass-artefacts and post-translational modifications (PTMs) as amino acid misincorporations (See Methods for details and Figure 1c). We identified a large number of modified peptides but only considered those which were also identified as un-modified and were uniquely assigned to a protein.

We identified at least one amino acid substitution in 180 datasets of the 283 *S. cerevisiae* datasets, we obtained from PRIDE (Jones and Côté, 2008; Perez-Riverol et al., 2018, 2022). The identification of amino acid misincorporation in ∼63% of examined dataset highlights that amino acid misincorporation are widespread and can be found in many existing massspectrometry datasets. We identified between 1 and 6583 amino acid misincorporations per dataset, totaling 47,917 amino acid misincorporations (Figure S1a). In order to highlight the generality of our approach, we also applied eTEL to 188 *E. coli* datasets from the PRIDE repository. We identified about twice as many amino acid misincorporation (91,565) in *E. coli* as in *S. cerevisiae*, ranging from 1 to 42,383 amino acid misincorporation per dataset (Figure S1b).

The number of detected amino acid misincorporation in *S. cerevisiae* and *E. coli* datasets is well correlated (two-sided Wald-test, Pearson’s *R*^2^ = 0.79, *p* < 10^− 30^, *n* = 180; Figure S1c, Pearson’s *R*^2^ = 0.72, *p* = 1.6 ×^− 16^, *n* = 94; Figure S1d) with the number of identified peptides on the logarithmic scale. This indicates that the number of identifiable amino acid misincorporations highly depends on the depth of a given mass-spectrometry dataset. While we observe twice as many amino acid misincorporation in *E. coli*, the number of identified peptides overall grows proportionally and as such, the error detection rate for the two organisms is on par. Specifically, the codon specific error detection rate for *S. cerevisiae* ranges from 1.51 × 10^−5^ to 4.8 × 10^−3^, while it ranges from 2.61 × 10^−5^ to 6 × 10^−3^ in *E. coli*.

### Nucleotide interactions determine codon/anticodon specific affinities

In order to understand differential amino acid misincorporation probabilities we need to model the mechanism underlying tRNA misincorporation. We modeled the codon/anticodon binding affinity as a unit-less parameter, describing not only binding strength but absorbing any additional linear effects of tRNA recognition by the ribosome, such as induced conformation changes and kinetic proof-reading. We assumed that the incorporation of an amino acid follows a multinomial distribution, where the incorporation probability is a function of tRNA abundance and codon/anticodon binding affinity. We fitted the nucleotide and position specific binding parameters to the observed amino acid misincorporation using a Markov Chain Monte Carlo (MCMC) algorithm with Gibbs sampling using a uniform prior distribution for all parameter. Each of the datasets was separately considered during the fitting process. Since the datasets used for model fitting are highly variable, we estimated parameter uncertainty due to dataset choice via bootstrapping. Datasets were bootstrapped 10,000 times and all parameter posterior means fell within the bootstrapped distributions (Table S1, Figure S2d,e).

As expected, fitted binding affinities show an increased affinity for the canonical Watson-Crick nucleotide pairs (Figure 1d). However, our fitted affinity matrix is not symmetric. For example, the codon/anticodon interaction U:G is favored, while the reverse interaction, G:U, is not (Figure 1d). The pyrimidine codon/anticodon pairs U:C, C:C, and U:U show the weakest binding affinities. Our nucleotide and position specific binding affinities allow estimating codon/anticodon binding affinities and disentangling the contributions of individual nucleotides at different positions.

The position-specific scaling term clearly disfavors nucleotide mismatches at the first and second position over the third (Figure S2a). Mismatches at the second codon position are the most unfavorable in *S. cerevisiae*, potentially caused by the steric issues such a mismatch would lead to, if the two flanking nucleotides are bound. In *E. coli*, however, mismatches at the first position are most unfavorable (Figure S2a). Watson-Crick pairings are always favorable, while most other nucleotide interactions are considered unfavorable (Figure S2b) for both *S. cerevisiae* and *E. coli*. Overall, the parameter sets correlate well (two-sided Waldtest, Pearson’s *ρ* = 0.91, *p* = 1.2 × 10^−7^, *n* = 18, Figure S2c) between the two species, likely due to the high degree of conservation of the translation machinery across the tree of life. However, both species differ significantly in all estimated parameters (Figure S2d), most likely because of differences in tRNA abundances.

### Translation error landscape is rugged

A wide variety of tRNAs can be wrongly incorporated at any codon. To describe the translation error landscape at the codon level, detected amino acid misincorporation across all *S. cerevisiae* datasets were pooled and plotted as a codon to amino acid misincorporation matrix (Figure 2a). Each entry represents the number of times a codon was observed to be wrongly translated with a given amino acid. Leucine and Isoleucine are treated as the same error because they cannot be distinguished in mass-spectrometry data due to their identical masses. We can clearly identify hotspots of substitutions, such as GAA (Glutamine acid) to Histidine or GUA (Valine) to Alanine. While the former requires two mismatches at the first and third codon position, the later requires one nucleotide mismatch at the second codon position. This is in contrast to *E. coli* in which these errors are underrepresented compared to *S. cerevisiae*. However, in *E. coli* there are other hotspots such as CUG and CUU to Valine substitution (Figure 2b, Figure S1e). Generally, we observe most mismatches at the first codon position, and a similar number of mismatches at the second and third codon position (41,924; 32,756; and 34,957 nucleotide mismatches), albeit seeing different types of nucleotide mismatches (Figure 2c). For example, G:G mismatches are most common at the first, C:U mismatches at the second codon position, and A:G mismatches are most common at the third codon position. However, if we consider only misincorporations that are best explained by single nucleotide mispairings, we find that mispairings at the first and second codon position cause the majority of tRNA mis-bindings (Figure 2d). Similar, the variability in the amino acids wrongly incorporated at a given codon also differs greatly between codons. For example, the Alanine codon GCG shows the least amount of variation in the misincorporated amino acids despite being the most error prone Alanine codon. In contrast, GCU appears to have the highest variance. This indicates that a codon’s error rate cannot be equated to the variation in amino acid misincorporations observed.

**Figure 2:**
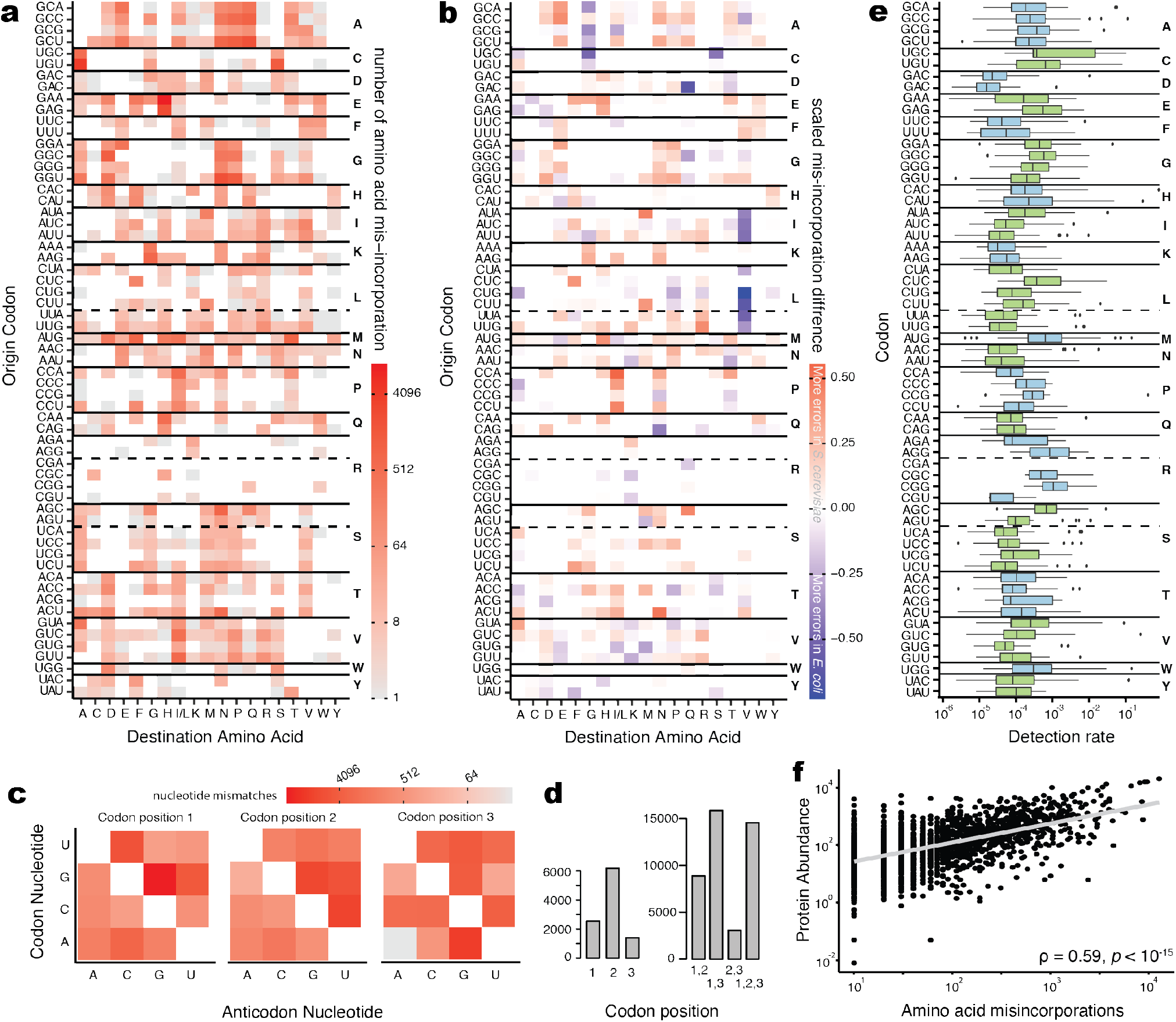
Codon choice determines translation errors. **a)** Heatmap of codon specific amino acid misincorporations for each target amino acid identified in the 180 datasets of *S. cerevisiae*. Synonymous codons often show the same errors (e.g., GCN to N, P, and Q), however many errors are detected only at individual codons (e.g., CGG to W). **b)** The translation error landscape of *E. coli* and *S. cerevisiae* differs substantially. Differences were calculated as the difference of the logarithm (augmented by one pseudo count) of the counts. **c)** Observed mistranslation events at individual codon sites. **d)** Multiple mispairings account for the majority of observed mistranslation events. However, when we consider only mispairing that are most likely caused by a single nucleotide mismatch, we find that position one and two show 2 and 3 times increase in mispairings over the third codon position, respectively. e) Codon-specific error detection rates for all codons vary greatly between synonymous codons, indicating large differences in fidelity between codons. Similarly, a large variation in the detection rate between datasets highlights the intrinsic noise in massspectrometry data. **f)** Detection of misincorporated amino acids correlates well with the integrated protein abundance for *S. cerevisiae*(Wang et al., 2015) (see also Figure S1).

### Codon-specific error detection rates vary by orders of magnitude

We calculated the error rate of a codon *i* by dividing the number of times codon *i* was observed to be mis-translated by the total number of times codon *i* was observed in a peptide (Figure 2e). The dependency of our error rate on the number and composition of the identified peptides likely results in an underestimate, as we are bound to miss the rarest of error events. We also observe a clear bias towards highly abundant proteins showing an increased number of amino acid misincorporations (ρ = 0.59, *p* < 10^−15^ Figure 2f). Amino acid misincorporations in low-abundance proteins are generally underrepresented in all the examined datasets. Due to these biases in detection, we will use the term error detection rate. Error detection rates vary by orders of magnitude between datasets and codons (Figure 2e). The Histidine codon CAU and the Tryptophan codon UGG show the highest variation among datasets ranging over three orders of magnitude, from 10^−5^ to 10^−2^. The Arginine codon CGA, one the other hand, was never detected to be substituted in *S. cerevisiae*. This might be explained by the rarity of CGA, which only represents 0.31 % of codons in the *S. cerevisiae* proteome (Tsuji et al., 2010). However, the lack of amino acid misincorporation observed at CGA codons could also be explained by the mis-translation of CGA as a STOP codon, producing peptides to which eTEL is currently blind. In *E. coli*, the Arginine codon CGA has a low error detection error rate of 6.03 × 10^−4^ (Figure S1f) and is likely detectable due to the increase in the overall number of detected amino acid misincorporations. In general, the six Arginine codons show less variation between datasets and a large variation in median error detection rate between codons, spanning two orders of magnitude, ranging from 1.95 × 10^−5^ for CGU to 1.05 × 10^−3^ for CGG (two-sided Wilcox rank sum-test ρ = 0.11; see Table S2 and Table S3 for all p-values between error rate pairs and sample sizes of *S. cerevisiae* and *E. coli*, respectively). The ability to quantify the variation in amino acid misincorporation rates is a clear advantage of proteomics approaches over rate quantification via individual constructs.

### Error free translation probability determined by codon composition

We explored how amino acid misincorporation are distributed across the proteome in order to better understand the effects of translation errors on individual proteins. We calculated the incorporation probability (*p*_*i*_) of each tRNA to each codon based on available tRNA abundance data from Weinberg et al., 2016 and based on our estimates of codon/anticodon binding affinities. The probability of incorporating a synonymous tRNA can then be calculated as

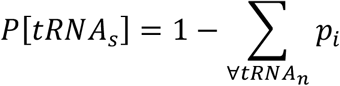

where *tRNA*_s_ is the set of synonymous tRNAs and *tRNA*_n_ is the set of non-synonymous tRNAs. We then mapped the probability of error free translation of a codon (*P*[*tRNA*_*s*_]) onto the proteome to investigate which proteins are most likely to be translated error free, or contain error hotspots.

When considering the whole proteome, amino acid misincorporations appear to be mostly uniformly distributed along the sequence of the proteins in *S. cerevisiae* (Figure S3a) and *E. coli* (Figure S3b). However, this does not hold true when considering individual proteins. The explicit site independence assumption of our model allows us to calculate the probability of error free translation until codon *i* as the cumulative product of error free translation probabilities until codon *i*. The cumulative error free translation probability for every individual protein reveals that the uniform distribution of amino acid misincorporations observed in the mass-spectrometry data is likely caused by differences in error probability along the codon sequence between proteins (Figure 3a). We find three general classes of error structures: i) slow initial decline in error-free probability with a high error probability towards the end of the protein (YLR249W, blue), ii) a constant decline in error-free probability (YBL104C, cyan), and iii) a stark initial decline in error-free probability and high fidelity towards the end of the protein (YOR127W, red). It is unclear if the error structure of a given protein is driven by selection or simply by chance.

**Figure 3:**
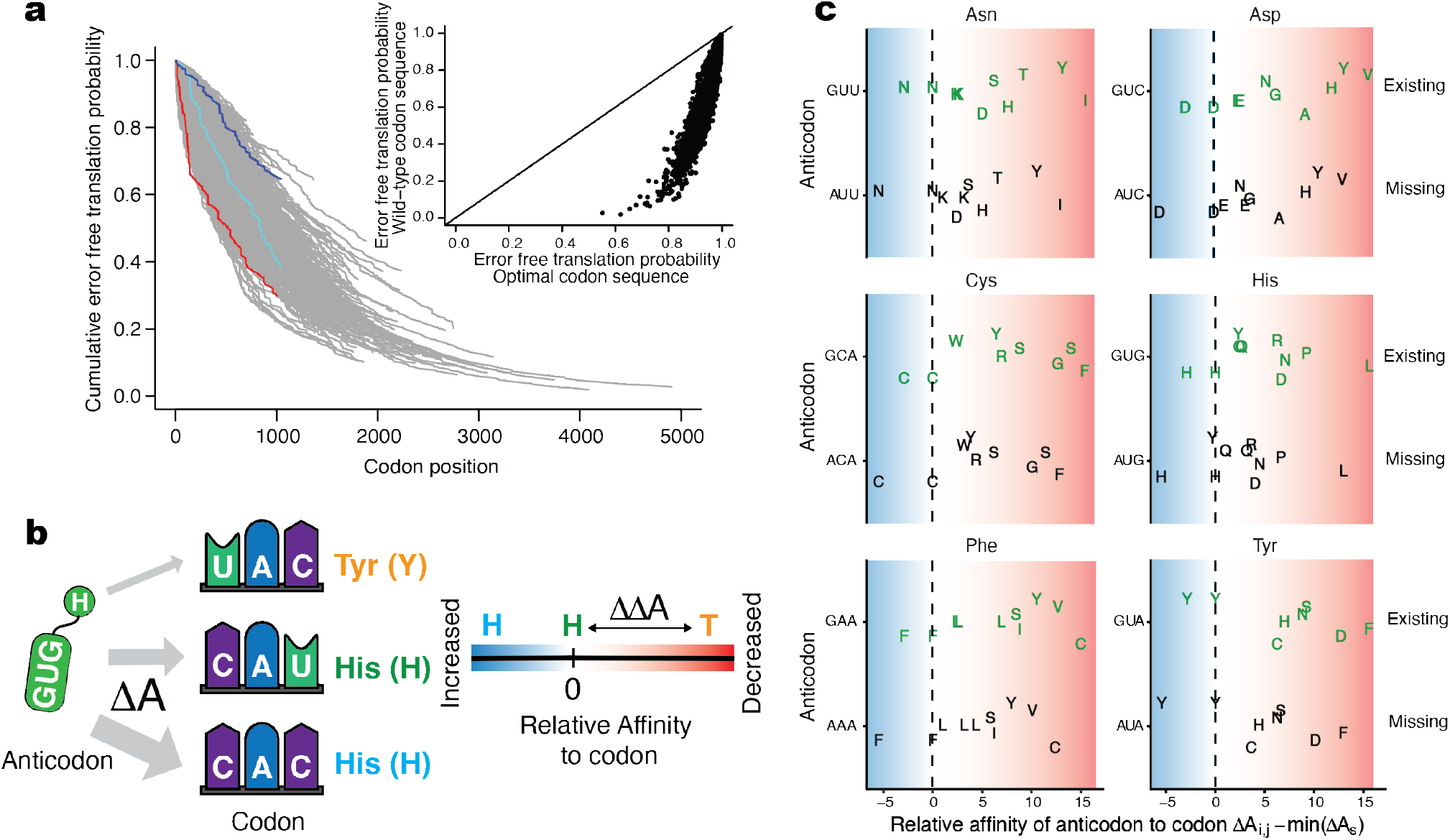
Codon choice and tRNA pool affect translation fidelity. **a)** Cumulative probability of error free translation for each protein in the S. cerevisiae proteome. Proteins exhibit different error probabilities along the sequence. Highlighted are three proteins with different error structures: i) increased error rates late (e.g., YLR249W, blue), ii) uniform error rates (e.g., YBL104C, cyan), and iii) increased error rates early (e.g., YOR127W, red) during translation. The inlay shows a comparison between the wild-type sequence error free translation probability and the hypothetical optimal sequences, that would minimize translation error probability. **b)** *T*o minimize mis-translations, non-synonymous codons should have a significantly lower binding affinity than synonymous codons. Relative binding affinity is quantified based on the lowest synonymous binding affinity. **c)** Impact of the tRNA pool on translation errors. Difference in codon/anticodon affinity for interactions with up to one codon/anticodon mismatch, for all two-codon amino acids with only one tRNA present in S. cerevisiae. The ribosome’s ability to discriminate between correct and incorrect binding events is diminished if we assume that the missing tRNA (black) would be present instead of the *existing* tRNA (green) (see also Figure S3).

### Translation efficiency and fidelity are not mutually exclusive

High variation in detection error rate between synonymous codons and their non-uniform occurrence across individual proteins led us to question the relationship between error rate and codon optimality. For example, the median error detection rate between arginine codons differs by almost two orders of magnitude, from 1.95 × 10^−3^ for AGG to 1.05 × 10^−3^ for CGG. We generally observe that a drastic increase in the probability of error free translation for most genes would be possible (Figure 3a inset) by using codons that have maximum fidelity. When we compare the codon adaptation index (Sharp and Li, 1987) (CAI) of wildtype *S. cerevisiae* codon sequences with the CAI of each sequence optimized for maximal error free translation we see drastic differences in their adaptiveness. In general, we see that CAI is a poor predictor for error free translation (Figure S3c). The increase in translation fidelity reduced the CAI for all codon adapted genes while increasing it for low expression genes which tend to be driven by mutation bias (Figure S3d). This indicates that the codon adaptation described by CAI does not reflect translation fidelity. Indeed, the relative synonymous codon usage is negatively correlated with the relative error detection rate (twosided Wald-test, *R*^2^ = 0.19, ρ = 0.0068, *n* = 58; Figure S3e), indicating that frequently used codons show higher translation fidelity. This relationship was previously observed(Mordret et al., 2019) and later confirmed by a more detailed analysis (Sun and Zhang, 2022), in *E. coli*. To further explore this trade-off, we compared our detection rates to estimates of ROCSEMPPR’s (Gilchrist et al., 2015) Δ*η*, which describes a codons translation inefficiency. Δ*η* decrease with decreasing inefficiency, meaning that the most efficient codons have the lowest value. Our observed error detection rates correlate well (two-sided Wald-test, *R*^2^ = 0.51, *p* = 3.5 × 10^−5^, *n* = 58, Figure S3f) with Δ*η* for *S. cerevisiae* taken from (Landerer et al., 2020), further indicating that there is no trade-off between translation efficiency and translation fidelity in *S. cerevisiae*. Thus, the reduction in CAI when optimizing a codon sequence for translation fidelity is not expected to be related to a decrease in translation efficiency. Providing further evidence that selection on codon usage is - at least in part - the result of selection for increased translation accuracy (Akashi, 1994).

### Absence of cognate tRNAs reduces amino acid misincorporations

No organism carries matching tRNA species for all codons. The absence of some tRNAs potentially serves to reduce ambiguity between tRNAs encoding different amino acids or to regulate translation speed (Ehrlich et al., 2021). Of the 61 × 61 = 3721 theoretical codon/anticodon interactions, only 2501 (67%) are possible in *S. cerevisiae*. The remaining 1220 interaction are impossible due to the lack of the corresponding tRNA. We tested if our affinity parameters differ between tRNA species present and absent. We investigated the difference in binding affinity of a given anticodon with its cognate and near-cognate codons (Figure 3b). For example, the Phe-tRNA^GAA^ has a reduced binding affinity towards the near-cognate Leucine codons UUA and UUG compared to its cognate UUC (Shah and Gilchrist, 2010). The affinity of the Phe-tRNA^GAA^ to UUA and UUG is only 77%, and 78% of the affinity to UUC, respectively. However, if we assume the Phe-tRNA would carry the AAA anticodon instead, the affinity of the Phe-tRNA towards UUA and UUG comes close to that of UUC (95% and 84%, respectively). Similar effects can be observed for other amino acids encoded by exactly two codons in *S. cerevisiae* and *E. coli* (Figure 3c; Figure S3g). It is, therefore, possible that the presence of tRNA species in both organisms has been under selection to minimize amino acid misincorporation by maximizing the affinity difference between synonymous and non-synonymous codons.

### Translation machinery is nearly optimal

Most proteins can be expected to carry at least one amino acid misincorporation. It is, therefore, important to understand the impact amino acid misincorporation have on protein function and fitness. We performed a proteome-wide estimation of fitness effects for all possible mutations at a given site using EVmutations (Hopf et al., 2017). For *S. cerevisiae* and *E. coli*, respectively, we calculated in total 26,504,184 and 13,117,068 fitness effects across 4584 and 3081 proteins with sufficient alignment depth. We combined our fitness estimates with amino acid misincorporation probabilities from mTEL and protein abundance from PAXdb (Wang et al., 2015) to understand fitness effects of amino acid misincorporations relative to an error free protein population (Figure 4a). We found that the severity of the calculated fitness effects declines with increasing probability of error occurrence in both *S. cerevisiae* (Figure 4b) and *E. coli* (Figure 4c), indicating potential selection on codon usage to avoid harmful amino acid misincorporations.

**Figure 4:**
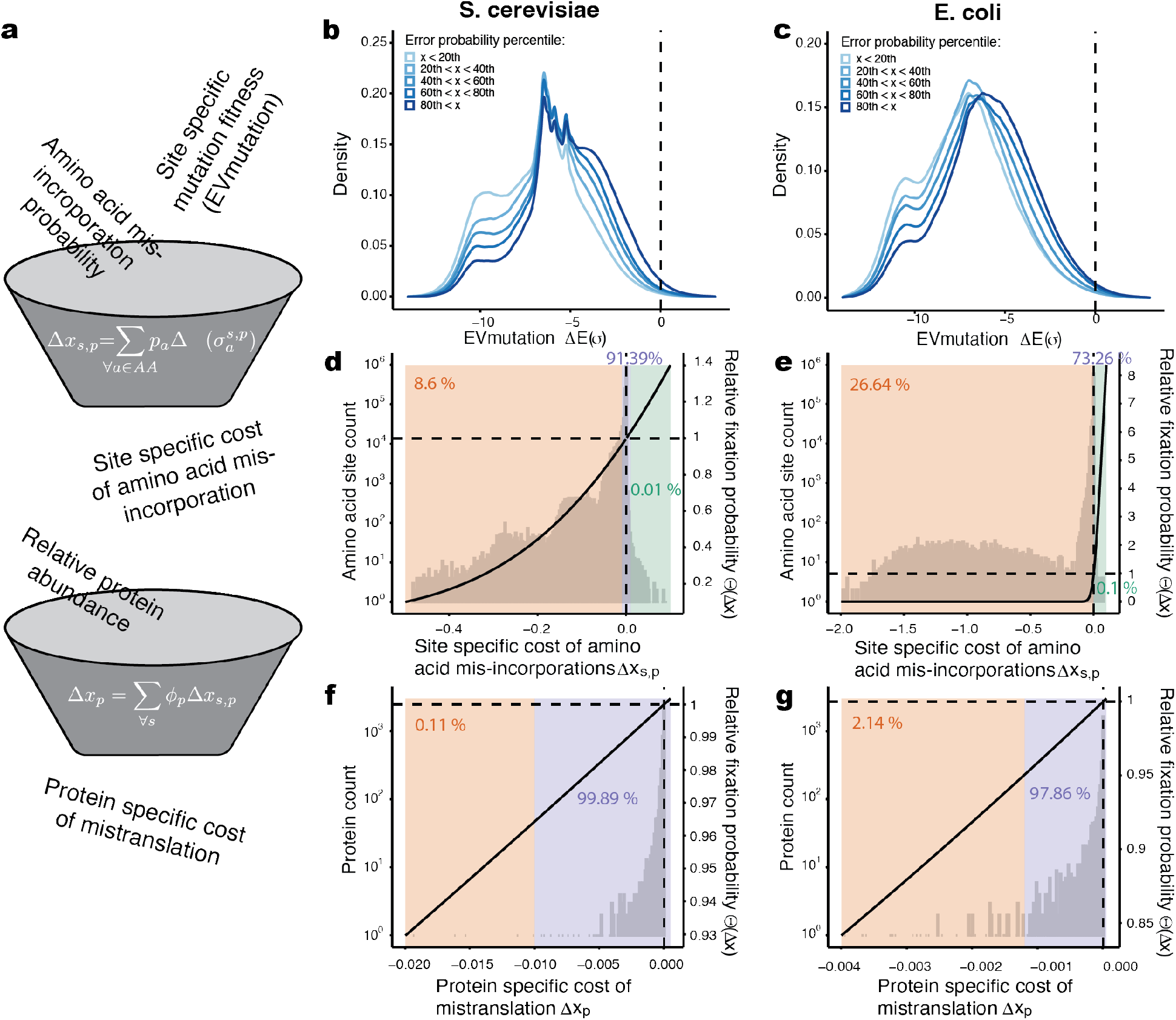
Deleterious amino acid misincorporations are compensated by reduced error rate. **a)** Site-specific fitness effects of amino acid mutations (EVmutations(Hopf et al., 2017)) were combined with amino acid misincorporations probabilities (mTEL) to calculate a site-specific expected cost of an amino acid misincorporation. Accounting for individual protein abundance and a protein’s contribution to the overall proteome, we can estimate the fitness cost of a protein due to mistranslation. **b, c)** The translation machinery prevents strongly deleterious misincorporations while allowing for advantageous ones. Severity of deleterious effects of substitutions declines with increased error probability (from light to dark blue). **d, e)** The majority of amino acid misincorporations are neutral (purple region), a smaller percentage (∼8.6 % in *S. cerevisiae* and ∼26.64 % in *E. coli*) are deleterious (orange region), and only a very small fraction is considered advantageous (green region). The black curve shows the relative fixation probability, and the dashed lines indicate the shift from decreased to increased relative fixation probability. **f, g)** Protein burden was calculated as the sum of individual site contributions and weighted by the relative contribution of a protein to the proteome based on protein abundance(Wang et al., 2015). Only 0.11 % of proteins impose a significant burden due to amino acid misincorporations in *S. cerevisiae* (**f**) while 2.14 % of proteins in *E. coli* (**g**).

These fitness effects assume that a mutation is present in 100% of the proteins. A translation error, however, will only affect the one protein that contains it. Combining the error probabilities with the calculated fitness effects, we can assess the expected effects amino acid misincorporations have on a protein. We calculated the fitness difference between the wildtype protein assuming perfect – error free – translation, and the effect an amino acid misincorporation has on the protein (Figure 4d,e). Assessing the relative fixation probability of amino acid misincorporations shows that the vast majority of amino acid misincorporations are neutral in both *S. cerevisiae* and *E. coli* (91.39% and 73.26%). In contrast, only a small number of amino acid misincorporations are deleterious (8.6% and 26.64%), or even advantageous (0.01% and 0.1%). This indicates that *S. cerevisiae* faces little to no selection to further invest into increasing translation fidelity and improve energetically expensive kinetic proofreading capabilities. *E. coli* appears to face much larger selection than *S. cerevisiae* on individual sites as, at almost one third of sites, amino acid misincorporations are deleterious.

### Maximizing translation fidelity is deleterious

We calculated the fitness effects of translation errors of individual proteins in *S. cerevisiae* and *E. coli* by summing the site-specific fitness effects. Protein fitness effects were then weighted by their protein abundance from PAXdb (Wang et al., 2015) and contribution to the total amount of protein produced. Only a small fraction of proteins impose a significant fitness burden on *S. cerevisiae* (∼0.11 %, Figure 4f) and on *E. coli* (∼2.14 %, Figure 4g) due to error prone translation. However, the summed fitness burden of all proteins is significant in both organisms (*S. cerevisiae*: Δ*x*_*p*_ = −0.789; *E. coli*: Δ*x*_*p*_ = −0.414). While the fitness burden in *E. coli* is less than in *S. cerevisiae*, the efficacy of selection is greater, with a relative fixation probability of ΘLΔ*x*_*p*_M = 0.019 in *S. cerevisiae* and ΘLΔ*x*_*p*_M = 3 × 10^−14^ in *E. coli*. This results in the fixation probability in both species to be drastically reduced relative to a hypothetical error free translation system. However, this reduction in fitness is conferred by only a small number of proteins in both species (Figure 4f,g) and therefore local solutions to increase fidelity may be preferred (Rajon and Masel, 2011). If we assume that there is no selection to further improve fidelity of the translation machinery we can estimate the fitness cost of global increases in translation fidelity e.g., by improved kinetic proofreading. The fitness cost imposed by an error free translation machinery would be 0.7753 and 0.41267 for *S. cerevisiae* and *E. coli*, respectively. This burden is on par with the cost of translation errors, making it unlikely that such an increase would reach fixation.

## Discussion

Detection and modeling of amino acid misincorporations at proteome-scale enabled us to estimate their fitness effects and to gain insights into the evolution of translation fidelity. Our mechanistic model of amino acid misincorporations (mTEL) was trained on misincorporations detected in existing mass-spectrometry data sets (eTEL). Using mass spectrometry, peptides can be detected and identified by matching measured peptide spectra to theoretical spectra generated from a sequence database (Cox and Mann, 2011). Many spectra, however, are not matched to any theoretical peptides, due to e.g., post-translational modifications (PTM) which alter the peptides’ mass. Various approaches have been developed to account for such mass-shifts due to PTMs (Chick et al., 2015; Tang et al., 2005; Yu et al., 2020). Mordret et al. were first to note that amino acid misincorporations in a peptide yield a similar mass shift and that strategies developed for the detection of PTMs can be used to identify amino acid misincorporation proteome-wide via mass-spectrometry (Mordret et al., 2019). Based on this approach, we developed eTEL, a high-throughput pipeline that can process hundreds of datasets and after quality filters, report amino acid substitutions in an automated way. We applied it to all existing *S. cerevisiae* and *E. coli* datasets in the PRIDE repository. We identified 47,917 amino acid misincorporations in *S. cerevisiae* and 91,565 in *E. coli*, highlighting the vast amount of, yet, unexplored information in publicly available mass-spectrometry datasets. eTEL allows exploring amino acid misincorporations across different organisms and conditions that alter translation fidelity, such as aging or cancer (Pataskar et al., 2022).

While mass spectrometry is the only method to detect amino acid misincorporations proteome-wide, it has limitations. The proteome coverage varied between 0.05 – 82.3 % of all proteins in *S. cerevisiae* (0.07 – 58.5 % in *E. coli*). Since most proteins are only identified via a single peptide, the data cover only ∼4.5 % (∼10 % in *E. coli*) of all codon positions in total, leaving a large number of positions unexplored. This highlights the necessity to extrapolate error rates via mechanistic models. In addition, there is large variation in the number of times individual peptides are observed, ranging from 1 to 688 in *S. cerevisiae* (1 to 689 in *E. coli*). In shotgun proteomics, whether or not a peptide is detectable will depend on the protein, the peptide’s environment within the protein, and on the peptide sequence itself. Proteins are less likely to be present in the sample in sufficient amounts if they are only expressed under certain conditions, challenging for sample preparation (e.g., membrane proteins), or have a low copy number (e.g., transcription factors). On the level of individual peptides, the amino acid composition strongly influences the observability. Factors like hydrophobicity, the ability to hold positive charges and the isoelectric point are predictors of whether a peptide will be detected or not (Cech and Enke, 2000; Fusaro et al., 2009).

Overall, the coverage of protein sequences by mass spectrometry is biased, and influenced by the properties of the corresponding peptides that are measured.

We focused on tRNA misincorporation as the main mechanism of amino acid misincorporation. However, amino acid misincorporations can be the result of tRNA mischarging, when the tRNA is charged by the wrong amino acid. The enzymes responsible for charging the tRNAs, amino-acyl tRNA-synthetases (Ling et al., 2009), have crossreactivity to similar amino acids, e.g. Ile-aa-tRNA synthase bind Leu, Val and Norvaline albeit with lower affinity (Bilus et al., 2019). We cannot distinguish if a detected amino acid misincorporation is the result of mischarging or binding of incorrect tRNA.

The correct tRNA incorporation depends on a multitude of factors such as codon/anticodon binding affinity (Gromadski and Rodnina, 2004; Pape et al., 2000), tRNA abundance (Sipley and Goldman, 1993), or the kinetic proofreading performed by the ribosome (Gromadski and Rodnina, 2004) (see (Rodnina et al., 2005) for review). We focus on tRNA abundance, which is readily available for a multitude of organisms, and codon/anticodon binding which can be parameterized with few, but meaningful parameters. Most of these factors controlling successful tRNA incorporation alone are too complex to be modeled in a meaningful manner, let alone interactions between them. mTEL allows us to describe the observed amino acid misincorporation and can be used for *in silico* experiments not feasible in the laboratory. For example, we tested the impact of missing tRNAs (Figure 3).

While our mechanistic model of ribosome/tRNA interactions (mTEL) was fitted to observed amino acid misincorporations, it can also predict probabilities of amino acid misincorporations not observed by mass spectrometry. For example, by leveraging information across codon sites and nucleotide interactions allowing us to estimate amino acid misincorporation at the Tryptophan codon without having it observed in *S. cerevisiae*.

Population genetics would predict that costly translation errors, which render a protein nonfunctional, should appear, if at all, early during protein synthesis in order to minimize wasted production costs. While this is plausible for non-sense errors such as premature termination or frameshifts, missense errors during early and late stages of protein synthesis incur the same production costs. In agreement with this, we do not observe any proteome-wide bias in the distribution of amino acid misincorporations along the protein sequence in *S. cerevisiae* (Figure S3a) and *E. coli* (Figure S3b).

We combined our predicted amino acid misincorporation probabilities with fitness estimates from EVmutation (Hopf et al., 2017) to describe the burden mis-translation events have on a cell. We accounted for the contribution of a protein to the overall protein pool by considering its abundance. However, we cannot consider the cost of a translational machinery with increased fidelity. It is therefore possible that the increased energy required to improve translation fidelity (‘global’ solution) exceeds the energy lost due to translation errors. Only about 8.6 % of codon sites appear to impose a significant fitness burden. Thus, ‘local’ adaptation (Rajon and Masel, 2011) of individual protein sequences, e.g., via codon choice, may be the preferred mechanism to cope with the remaining fitness burden imposed by mistranslations. Based on transcription error estimates, it was previously proposed that ‘local’ solutions on a gene-by-gene basis can reduce the effects of errors (Rajon and Masel, 2011). Here, based on data of proteome-wide translation errors, we detect ‘local’ solutions to amino acid misincorporations such as codon choice, and describe how ‘global’ solutions such as presence/absence of tRNA species can increase translation fidelity.

Amino acid misincorporations are only one type of synthesis errors introduced by the translation step. Other errors include frameshifts by ribosomal slippage, premature termination by falling of the ribosome, and stop codon readthrough by amino acid misincorporation or skipping. The evolutionary impact of phenotypic mutations in general is still unclear. They contribute to protein diversity and can have adaptive function, e.g., frameshifts in viruses, bacteria and yeast (Romero Romero et al., 2022). Yet, most phenotypic mutations are likely stochastic events, errors without known functions (Li and Zhang, 2019).

There is a large variability in translation fidelity across organisms based on individual reporter assays (Romero Romero et al., 2022). Using deTEL, it is now possible to explore translation accuracy in many species as well as under changing conditions, such as stress (Mohler and Ibba, 2017) or diseases (Kirchner and Ignatova, 2015) like cancer (Goodarzi et al., 2016; Pataskar et al., 2022). Although phenotypic mutations are individually rare, collectively, they can exert a significant fitness effect, and play a critical role in evolution.

## Acknowledgements

This work was funded by the Max Planck Society MPRGL funding. We thank HongKee Moon from the Scientific Computing Facility for his support in software engineering. We thank Michele Marass and Andrej Shevchenko for the critical reading of the manuscript. We thank Federica Luppino for her support with EVcouplings.

## Author Contribution

Conceptualization: A.T.P., and C.L. Data curation: C.L. Formal analysis: C.L Funding acquisition: A.T.P Investigation: A.T.P, and C.L. Methodology: C.L, and J.P Project administration: A.T.P Resources: A.T.P Software: C.L, and J.P Visualization: C.L Writing–original draft: A.T.P, and C.L Writing–review & editing: A.T.P, C.L, and J.P

## Code and data availability

The deTEL pipeline developed and used as part of this manuscript is available at https://git.mpi-cbg.de/tothpetroczylab/deTEL.

## Methods

A Full description of the Materials and methods can be found in the supplementary materials.

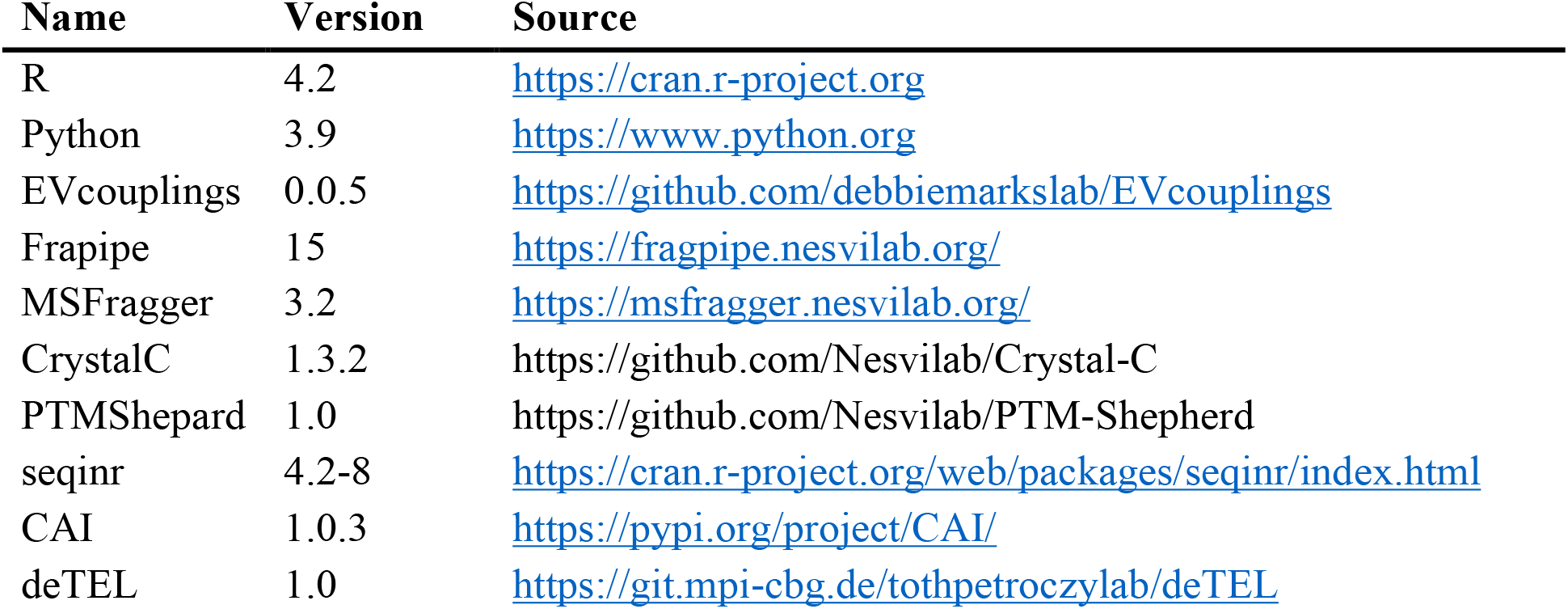

### Detecting amino acid misincorporations via open search

Datasets downloaded from PRIDE(Jones and Côté, 2008; Perez-Riverol et al., 2018, 2022) (last accessed: 04.11.2021) were analyzed in bulk by first using the dry-run functionality of the Fragpipe (version 17) software suite to extract the commands executed by Fragpipe. The individual tools included in Fragpipe were then called from a bash script on a computing cluster, searching each PRIDE project in parallel. Options are based on the settings used in the predefined ‘Open Search’ of Fragpipe. All raw files (Thermo .raw format) in the dataset were then searched together by MSFragger(Kong et al., 2017) (version 3.4) open search(Yu et al., 2020) against the *S. cerevisiae* proteome (translated reference CDS from yeastgenome.org, last accessed 12.03.2020) with mitochondrial genes removed and the *E. coli* proteome (extracted from genome and translated, genome obtained from NCBI, last accessed 15.09.2020).

### Detecting substitutions in open search results

Amino acids substitutions were detected with a custom Python script (version 3.9). For each dataset, FDR-filtered psm.tsv files were collected. Peptides were further filtered to remove all that were matched to several proteins. All peptides were annotated with their start and end position. Peptides with a mass shift between −5 and +5 mDa were considered as unmodified, and all others as modified. Modified peptides were only retained if the modified position in the protein sequence was covered by an unmodified peptide present in the same MSFragger file (identified in the same MS measurement) and if the position of the modification could be unambiguously localized. All remaining peptides with a mass shift matching the mass difference between two amino acids were marked as substitutions, and the original and substituted amino acid were annotated, with Leucine and Isoleucine treated as equivalent. All peptide with a mass shift and localization also matching a known PTM (see supplementary data for a full list) were removed. For each codon, a detection rate was calculated as the total number of times (number of PSMs) a peptide covering a position with this codon and carrying a substitution at that position was detected divided by the number of times the same codon was detected on a peptide unmodified.

### Exploring the relationship between codon usage and translation fidelity

Codon adaptation index (CAI) was calculated using the R (version 4.2.0) package seqinr (version 4.2-8) using the function CAI. Codons weights were taken from seqinr’s caitab. Relative synonymous codon usage was calculated from the top 5% expressed S. cerevisiae genes using gene expression (RPKM) data from(Weinberg et al., 2016) using the CAI (version 1.0.3) Python (version 3.9) package.

### Modeling error prone protein production

We designed a system of equations, modeling the production of proteins from DNA. As we are only interested in the production of proteins and not in their lifetime, we ignore degradation.

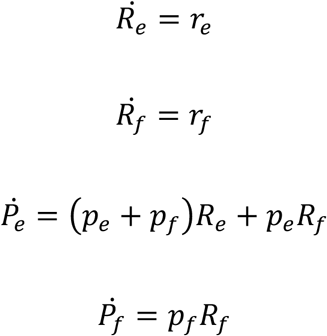

Integrating the system and solving it at time *t* = 1 allows us to determine the dependency of error prone protein (*P*_*e*_) produced by transcription and translation.

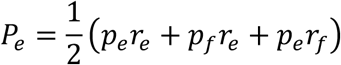

which we can separate into the contribution of transcription

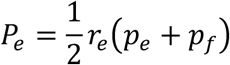

and translation

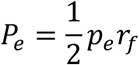

We assume further, that the total amount of mRNA and protein produced is *r*_*t*_ = *r*_*e*_ + *r*_*f*:_ and *p*_*t*_ = *p*_*e*_ + *p*_*f*_, respectively. This assumption allows us to compare literature estimates of misincorporation rates to our model.

### tRNA arrival probabilities

While we assume the binding probabilities of the tRNAs at stationarity, we have to consider the arrival rates of each tRNA at the ribosome. We assume that the waiting time for a tRNA to arrive at the ribosome is exponentially distributed with the rate being proportional to the tRNA abundance. tRNA abundance was obtained from Weinberg et al(Weinberg et al., 2016) and Larson et al(Larson et al., 2014) for *S. cerevisiae* and *E. coli*, respectively. The rate parameter for the exponential distribution is calculated following(Fluitt et al., 2007).

### tRNA binding probabilities

The binding probability of anticodon *j* to codon *i* depends on the binding affinity Δ*A*_*i,j*_. Binding is parameterized based on the individual nucleotide binding affinities 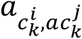 where 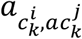 is the binding affinity between the nucleotide *c* of codon *i* at position *k* and the anticodon *j* nucleotide *ac* at position *k*. Binding affinities are further scaled by site specific importance terms *s*_*k*_.

### tRNA incorporation probability

To calculate the incorporation probability of a tRNA we have to distinguish two cases. First, the tRNA *a* in question arrived before any competitor. Second, a competitor arrived before. In the first case, it is simply that the probability to bins first is equal to the probability to bind, 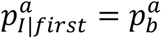, as nothing else is to consider. In the second case, however, we have to consider each tRNA that may have arrived at the ribosome ahead of the focal tRNA. Considering that every tRNA arriving before has to be rejected *λ*_*k*_/*λ*_*a*_ times to allow for the focal tRNA to bind, we find that

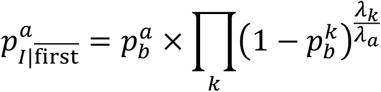

### Model fitting

We fitted the multinomial Translation Error Landscape (mTEL) model using Markov Chain Monte Carlo with Gibbs sampling and uniform priors on all parameters. The Markov chain was estimated for 10,000 steps after an initial burn-in period of 10,000 steps. Only every 10th step was retained and the last 500 thinned samples were used as posterior distribution.

### Calculating Fitness effects

EVcouplings (version 0.0.5) was used to create fitness estimates of amino acid substitutions. We collected all Uniprot IDs assigned to the *S. cerevisiae* and *E. coli* reference proteomes. The EVcouplings(Hopf et al., 2018) alignment step was performed for each protein with bit scores 0.1 – 0.5. The alignment with the best performing bit score was selected for the Fitness estimation. Fitness effects of all amino acid substitutions were estimated as Δ*E*(σ) using EVmutation(Hopf et al., 2017) based on co-evolution and conservation of residues. Sitespecific fitness effects Δ*x*_*s,p*_ of translation errors were then defined as the weighted average amino acid fitness at a given position. The evolutionary effect of each amino acid misincorporation was assessed following the fixation probability definition of Sella and Hirsh(Sella and Hirsh, 2005). The effective population size was assumed to be *N*_9_ = 8,600,000 in *S. cerevisiae*(Tsai et al., 2008) and *N*_*e*_ = 10^8^ in *E. coli*(Lynch, 2010). A scaling Parameter *q* = 4.19 × 10^−7^ represents the value of an ATP(Gilchrist, 2007). Protein specific fitness effects Δ*x*_*p*_ were calculated as the sum of the site-specific fitness effects Δ*x*_*s,p*_ weighted by the relative contribution of a protein to the proteome based on its abundance. Relative protein abundance ϕ_*p*_ of a protein was derived from the integrated data from PAXdb(Wang et al., 2015) for *S. cerevisiae* and *E. coli*, respectively. Protein specific evolutionary effects were assessed in the same way as site-specific effects.

### Statistical analysis and data visualization

was performed by R. For all box plot representations thick black line indicates median, box indicates 25^th^ and 75^th^ percentiles, whiskers indicate 1.5 times the inter-quartile range.

## Supplementary information titles and captions

**Figure S1: Amino acid misincorporations are abundant in many existing massspectrometry datasets. a)** Number of identified substitutions in the 180 S. cerevisiae datasets with identified amino acid misincorporations. **b)** Detected substitutions increase with identified peptides. Datasets where selected (blue dots) if they contained at least 10 substitutions were detected and where not considered outliers by our regression. **c)** Number of identified substitutions in the 94 E. coli datasets with identified amino acid misincorporations. **d)** Detected substitutions increase with identified peptides. Datasets where selected (blue dots) if they contained at least 10 substitutions were detected and where not considered outliers by our regression. **e)** Heatmap of all codon specific amino acid substitution identified across 94 datasets. Leucine and Isoleucine cannot be distinguished due to their identical mass. Each cell shows the number of misincorporations of amino acid ***i*** observed for a codon ***j***. In contrast to S. cerevisiae, we find the Tryptophan codon TGG misincorporated as a variety of amino acids. **f)** Codon specific error detection rate in E. coli. We observe similar variability of error rates between synonymous codons and datasets to S. cerevisiae. For most amino acids, E. coli and S. cerevisiae share the codon with the highest fidelity. Notable exceptions are Leucine and Valine (thick black line indicates median, box indicates 25^th^ and 75^th^ percentiles, whiskers indicate 1.5 times the inter-quartile range).

**Figure S2: Parameter bootstrap and model parameter posterior distributions between S. cerevisiae and E. coli. a)** Distribution of position posterior means. **b)** Distribution of wobble parameter posterior means. **a, b)** Dashed line indicates model fit. **c)** Distribution of the position posteriors. **d)** Distribution of wobble parameter posteriors. **c, d)** All distributions show a significant difference in their mean values between S. cerevisiae and E. coli (KruskalWallis test, *ρ* < 0.0001, *n* = 500).

**Figure S3: Exploring effects of codon/anticodon usage on translation efficiency fidelity**. **a)***S. cerevisiae* shows a mostly uniform distribution of amino acid misincorporations across all proteins in the proteome. **b)** *E. coli* also shows a mostly uniform distribution. In contrast to *S. cerevisiae*, however, a noticeable drop-off in detected misincorporations can be observed in the last bin; potentially explained by the lack of tryptic peptides at the end of a protein. **a, b)** Each bin represents one percent protein length. **c)** Codon adaptation index (CAI) is only weakly correlated with error free translation probability. While genes with high CAI are more likely to be translated error free, the opposite does not hold as many proteins with low CAI show high fidelity. **d)** Comparing the codon adaptation index (CAI) wild-type and fidelity optimized codon sequences reveals a drop in sequence adaptation for sequences with a high CAI. Wild-type sequences with a low CAI and a codon usage likely dominated by mutation tend to improve their codon adaptation. **e)** Relative synonymous codon usage describing the preferred codon usage of the most highly expressed genes (top 5 %) is negatively correlated with the relative codon detection rate indicating that frequent codons are translated more accurately. **f)** ROC-SEMPPR’s *Δη* values, describing a codons translation inefficiency (increasing *Δη* indicates inefficient codons) are positively correlated with a codons error detection rate. Indicating that inefficiently translated codons tend to have a higher error rate. **g)** Impact of the tRNA pool on translational errors. Difference in codon/anticodon affinity for interactions with up to one codon/anticodon mismatch, for all two-codon amino acids with only one tRNA present in *E. coli*. The ability of the ribosome to discriminate between correct and incorrect binding events is diminished if we assume that the missing tRNA (black) would be present rather than the tRNA naturally present (green).

**Table S1: Model parameter posterior means with empirical p-value based on 10.000 bootstrap samples for S. cerevisiae**. Empirical p-value was calculates as (r + 1)/(n + 1) were r is the number of sample greater than the posterior mean, and n is the total number of samples(Davison and Hinkley, 1997).

**Table S2: Comparison of error detection rate distributions for *S. cerevisiae***. Error rate distributions were compared using a two-sided Wilcox test. The upper triangle of the table indicates the significance level as p-values, while the lower table indicates the sample size for each distribution in the format <row>/<column>. See supplementary_table_2_3.xlsx

**Table S3: Comparison of error detection rate distributions *E. coli***. Error rate distributions were compared using a two-sided Wilcox test. The upper triangle of the table indicates the significance level as p-values, while the lower table indicates the sample size for each distribution in the format <row>/<column>. See supplementary_table_2_3.xlsx

### Data S1. (yeast_fitting.zip)

- **fitted_energies.csv:** Codon/Anticodon binding affinities for all possible interactions for *S. cerevisiae*.
- **fitted_substitution_probabilities.csv:** Fitted amino acid incorporation probabilities for each codon for *S. cerevisiae*.
- **likelihood_trace.csv:** Likelihood trace.
- **position_trace.csv:** Position parameter trace.
- **wobble_trace.csv:** Codon/Anticodon affinity trace.
- **position.csv:** Position parameter posterior distribution
- **wobble.csv:** Codon/Anticodon affinity parameter posterior distribution.

### Data S2. (yeast_data_summary.zip)

- **yeast_substitutions.zip:** Detected amino acid misincorporations, number of codons observed in error free and error prone peptides, and number of detected peptides per protein in *S. cerevisiae* datasets from PRIDE.
- **yeast_tRNA_count.csv:** tRNA abundance used for mTEL model fitting.
- **yeast_ppm_paxdb_integrated:** Protein abundance data used in Figure 4f.

### Data S3. (ecoli_fitting.zip)

- **fitted_energies.csv:** Codon/Anticodon binding affinities for all possible interactions for *E. coli*.
- **fitted_substitution_probabilities.csv:** Fitted amino acid incorporation probabilities for each codon for *E. coli*.
- **likelihood_trace.csv:** Likelihood trace.
- **position_trace.csv:** Position parameter trace.
- **wobble_trace.csv:** Codon/Anticodon affinity trace.
- **position.csv:** Position parameter posterior distribution
- **wobble.csv:** Codon/Anticodon affinity parameter posterior distribution.

### Data S4. (ecoli_data_summary.zip)

- **ecoli_pride.zip:** Detected amino acid misincorporations, number of codons observed in error free and error prone peptides, and number of detected peptides per protein in E. coli datasets from PRIDE.
- **ecoli_tRNA_count.csv:** tRNA abundance used for mTEL model fitting.
- **ecoli_ppm_paxdb_integrated:** Protein abundance data used in Figure 4g.

