## Supplementary Material for "Evolutionary impact of codon specific translation errors at the proteome scale"

<u>Detecting amino acid misincorporations via open search</u>

Datasets downloaded from PRIDE[26–28] (last accessed: 04.11.2021) were analyzed in bulk by first using the dry-run functionality of the Fragpipe (version 17) software suite to extract the commands executed by Fragpipe. The individual tools included in Fragpipe were then called from a bash script on a computing cluster, searching each PRIDE project in parallel. The configuration files and command line options are based on the settings used in the predefined 'Open Search' workflow in the GUI version of Fragpipe. For each dataset, a 'workspace' was first initialized by Philosopher. All raw files (Thermo .raw format) in the dataset were then searched together by MSFragger[30] (version 3.4) open search[31] against the *S. cerevisiae* proteome (translated reference CDS from yeastgenome.org, last accessed 12.03.2020) with mitochondrial genes removed and the *E. coli* proteome (extracted from genome and translated, genome obtained from NCBI, last accessed 15.09.2020).

MSFragger was set to perform mass calibration, deisotoping and to remove neutral losses. True precursor tolerance was 5 ppm, MS/MS tolerance was 10 ppm. The range of mass shifts (precursor mass window) was set to -135 to 135 Da, excluding the range from -1.5 to +3.5 Da. MSFragger was set to allow for any mass shift that fell within this range and localized to any amino acid. In-silico digestion was specified as 'strict trypsin' (cut after K and R, regardless of C-terminal amino acid), with up to two missed cleavages. Only fully tryptic peptides with a length of 7 – 50, a mass of 500 – 5000 Da, and a precursor charge of 1 – 4 were allowed. A fixed modification of Cystein with mass 57.02146 (carbamidomethylation), and no variable modifications. All other MSFragger settings were kept to the default. MSFragger output (.pepXML and .tsv files) were move to a dataset-specific folder and processed by CrystalC[65] (version 1.3.2) to identify missed cleavages, semi-enzymatic peptides and chimeric spectra among the MSFragger identifications. Precursor charge was set to 1 – 6, precursor mass tolerance to 20 ppm, and the precursor isolation window to 0.7 Da. Number of theoretical isotope peaks was set to 3, and isotope error correction was disabled.

PeptideProphet was then used to score the peptides retained by CrystalC. The c.pepXML files produced by CrystalC were analysed with the options –-nonparam, --decoy, --decoyprobs and –expectscore enabled, the mass width set to 1000 and the clevel set to 2. PeptideProphet[66] output was combined into a single .interact.pepXML file. ProteinProphet[66] was then executed on PeptideProphet's output to score proteins, with the maximum ppm difference set to 2,000,000 (standard in the Open Search) and combining the output. The Philosopher[66] commands 'annotate', 'filter', and 'report' were called to produce additional reports. 'annotate' was called on the sequence database with standard parameters. 'filter' was called on the .pepXML and .protXML files generated by PeptideProphet and ProteinProphet, respectively, with the –sequential, --razor, and –mapmods options enabled, and the protein-level FDR set to 0.01. 'report' was called with the —decoys option enabled. The workspace was cleaned up by the 'workspace' command. A summary of the identified mass shifts was generated with PTM-Shepherd[67] (version 1.0). All options were left at default, the only specified variable modification was failed carbamidomethylation with a mass of -57.021464.

<u>Detecting substitutions in open search results</u>

Amino acids substitutions were detected with a custom Python script (version 3.9). All generated and FDR filtered psm.tsv files were collected. Peptides were filtered to remove all that were matched to several proteins. All peptides were annotated with their start and end position. Peptides with a mass shift between -5 and +5 mDa were considered as unmodified, and all others as modified. PSMs associated with unmodified peptides were retained and all remaining decoys were removed.

Modified peptides were only retained if the unmodified peptide was present in the same MSFragger file (identified in the same MS measurement) and if the position of the modification could be unambiguously localized. Remaining peptides with a mass shift matching the mass difference between two amino acids were marked as substitutions, and the original and substituted amino acid were annotated, with Leucine and Isoleucine treated as equivalent. Additionally, all peptide with a mass shift and localization also matching a known PTM were removed.

Exploring the relationship between codon usage and translation fidelity

Codon adaptation index (CAI) was calculated using the R (version 4.2.0) package seqinr (version 4.2-8) using the function CAI. Codons weights were taken from seqinr's caitab. Relative synonymous codon usage was calculated from the top 5% expressed S. cerevisiae genes using gene expression (RPKM) data from[29] using the CAI (version 1.0.3) Python (version 3.9) package.

Modeling error prone protein production

We designed a system of equations, modeling the production of proteins from DNA. As we are only interested in the production of proteins and not in their lifetime, we ignore degradation.

$$\dot{R_e} = r_e$$
$$\dot{R_f} = r_f$$

$$\dot{P_e} = (p_e + p_f)R_e + p_e R_f$$

$$\dot{P_f} = p_f R_f$$

Integrating the system and solving it at time $t = 1$ allows us to determine the dependency of error prone protein ($P_e$) produced by transcription and translation.

$$P_e = \frac{1}{2}(p_e r_e + p_f r_e + p_e r_f)$$

which we can separate into the contribution of transcription

$$P_e = \frac{1}{2} r_e (p_e + p_f)$$

and translation

$$P_e = \frac{1}{2} p_e r_f$$

We assume further, that the total amount of mRNA and protein produced is $r_t = r_e + r_f$ and $p_t = p_e + p_f$, respectively. This assumption allows us to compare literature estimates of misincorporation rates to our model.

tRNA arrival probabilities

While we assume the binding probabilities of the tRNAs at stationarity, we have to consider the arrival rates of each tRNA at the ribosome. We assume that the waiting time for a tRNA to arrive at the ribosome is exponentially distributed with the rate being proportional to the tRNA abundance. tRNA abundance was obtained from Weinberg et al[29] and Larson et al[58] for *S. cerevisiae* and *E. coli*, respectively. The rate parameter for the exponential distribution is calculated following[24]. Briefly, we assume that a cell can be discretized into $n = V/l^3$ locations where $V$ is the cell volume (e.g. $0.6 \times 10^{-18} m^3$ for *E. coli* and

$4.2 \times 10^{-17} m^3$ for *S. cerevisiae*) and $l$ is the effective length of a tRNA ($1.58 \times 10^{-8} m$)[29]. Assuming a transition time of $\tau = l^2/6 * D$, where $D$ is the diffusion coefficient for tRNA ($8.42 \times 10^{-11} m^2$)[29] the arrival probability can be expressed as:

$$\lambda_i = \frac{P_i}{\tau}$$

were $P_i = [tRNA_i]/n$ is the probability of tRNA $i$ to occupy a given position in the cell.

Given the rate at which a tRNA arrives at the ribosome, we can calculate the probability that tRNA $a$ will arrive before tRNA $b$. We consider the joint probability of two tRNAs arriving at the ribosome.

$$f_{a,b} = f_{A|B(a|b)*f_B(b)}$$

$$\int_a \int_b f_{a,b} db\, da = 1$$

we can calculate the probability of $a$ arriving before $b$ as

$$p_{a<b} = \int_{a=0}^{\infty} \int_{b=a}^{\infty} f_{a,b} db\, da$$
$$= \int_{a=0}^{\infty} f_A(a) F_B(a) da$$
$$= \int_{a=0}^{\infty} \mu e^{-\mu a} \left(1 - e^{-\lambda a}\right) da$$
$$= \frac{\lambda}{\lambda + \mu}$$

Similarly, for many competitors, it holds that

$$\int_a \cdots \int_z f_{a,\cdots,z} dz \cdots da = 1$$

and we can express the probability of a focal tRNA $a$ to arrive before any other tRNA as

$$p_{a,\text{first}} = \frac{\lambda_a}{\sum_i \lambda_i}$$

tRNA binding probabilities

The binding probability of anticodon $j$ to codon $i$ depends on the binding affinity $\Delta A_{i,j}$. Binding is parameterized based on the individual nucleotide binding affinities $a_{c_k^i, ac_k^j}$ where $a_{c_k^i, ac_k^j}$ is the binding affinity between the nucleotide $c$ of codon $i$ at position $k$ and the anticodon $j$ nucleotide $ac$ at position $k$. Binding affinities are further scaled by site specific importance terms $s_k$. Thus, the binding affinity for each codon/anticodon pair $i, j$ can be calculates as

$$\Delta A_{i,j} = \sum_{k=1}^{3} a_{c_k^i, ac_k^j} s_k$$

In order to keep the system identifiable, we fix $s_3 = 1$. The Binding probability is then calculated as

$$p_{i,j}^b = \frac{\exp[-\Delta A_{i,j}]}{\sum_{\forall k} \exp[-\Delta A_{i,k}]}$$

where the denominator is the sum over the binding affinities for possible tRNAs.

<u>tRNA incorporation probability</u>

To calculate the incorporation probability of a tRNA we have to distinguish two cases. First, the tRNA $a$ in question arrived before any competitor. Second, a competitor arrived before.

Thus, according to the law for total probability

$$p_I^a = p_{I|\text{first}}^a \times p_{\text{first}}^a + p_{I|\overline{\text{first}}}^a \times p_{\overline{\text{first}}}^a$$

In the first case, it is simply $p_{I|first}^a = p_b^a$ as nothing else is to consider. In the second case, however, we have to consider each tRNA that may have arrived at the ribosome ahead of the focal tRNA. Considering that every tRNA arriving before has to be rejected to allow for the focal tRNA to bind, we find that

$$p_{I|\overline{\text{first}}}^a = p_b^a \times \prod_k \left(1 - p_b^k\right)^{\frac{\lambda_k}{\lambda_a}}$$

<u>Model fitting</u>

We fitted the multinomial Translation Error Landscape (mTEL) model using Markov Chain Monte Carlo with Gibbs sampling and uniform priors on all parameters. The Markov chain was estimated for 10,000 steps after an initial burn-in period of 10,000 steps. Only every 10 step was retained and the last 500 thinned samples were used as posterior distribution.

<u>Calculating fitness effects</u>

EVcouplings (version 0.0.5) was used to create fitness estimates of amino acid substitutions. We collected all Uniprot IDs assigned to the *S. cerevisiae* and *E. coli* reference proteomes. The EVcouplings[59] alignment step was performed for each protein with bit scores 0.1 – 0.5. The alignment with the best performing bit score was selected for the Fitness estimation. Fitness effects of all amino acid substitutions were estimated as $\Delta E(\sigma)$ using EVmutation[39] based on co-evolution and conservation of residues. Site specific fitness effects $\Delta x_{s,p}$ of translation errors were then defined as the weighted average amino acid fitness at a given position

$$\Delta x_{s,p} = \sum_{\forall a \in AA} p_a \, \Delta E\left(\sigma_a^{s,p}\right)$$

were $p_a$ is the probability of the amino acid misincorporation. The evolutionary effect of each amino acid misincorporation was assessed following the fixation probability definition of Sella and Hirsh[60]. The relative fixation probability of the observed fitness effects $\Delta x$ to a hypothetical error free translation system serving as a null hypothesis evaluated as

$$\Theta(\Delta x) = \frac{\big(1 - \exp(-2\Delta x q)\big) N_e}{1 - \exp(-2 N_e \Delta x q)}$$

where the effective population size is $N_e = 8{,}600{,}000$ in *S. cerevisiae*[61] and $N_e = 10^8$ in *E. coli*[62], $q = 4.19 \times 10^{-7}$ is the value of an ATP[63].

Protein specific fitness effects $\Delta x_p$were calculated as the sum of the site-specific fitness effects $\Delta x_{s,p}$ weighted by the relative contribution of a protein to the proteome based on its abundance.

$$\Delta x_p = \sum_{\forall s} \phi_p \, \Delta x_{s,p}$$

The relative protein abundance $\phi_p$ of a protein was derived from the integrated data from PAXdb[40] for *S. cerevisiae* and *E. coli*, respectively. Protein specific evolutionary effects were assessed in the same way as site-specific effects.

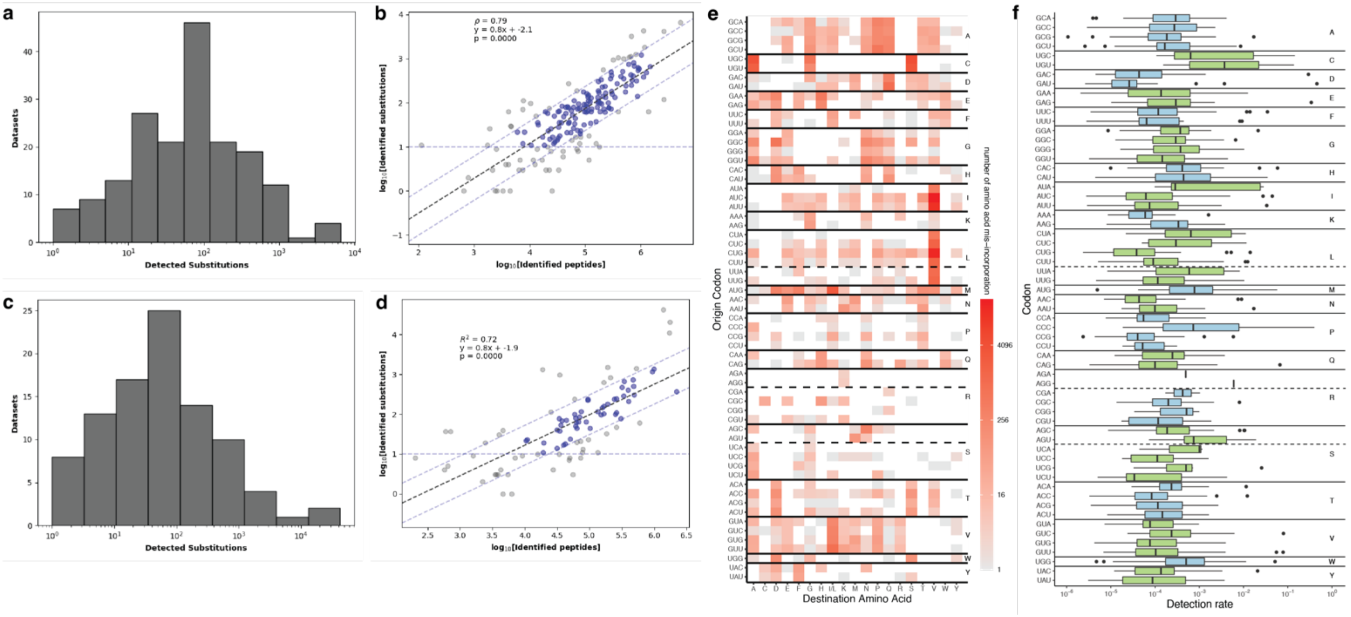


**Figure S1: Amino acid misincorporations are abundant in many existing mass-spectrometry datasets. a)** Number of identified substitutions in the 180 *S. cerevisiae* datasets with identified amino acid misincorporations. **b)** Detected substitutions increase with identified peptides. Datasets where selected (blue dots) if they contained at least 10 substitutions were detected and where not considered outliers by our regression. **c)** Number of identified substitutions in the 94 *E. coli* datasets with identified amino acid misincorporations. **d)** Detected substitutions increase with identified peptides. Datasets where selected (blue dots) if they contained at least 10 substitutions were detected and where not considered outliers by our regression. **e)** Heatmap of all codon specific amino acid substitution identified across 94 datasets. Leucine and Isoleucine cannot be distinguished due to their identical mass. Each cell shows the number of misincorporations of amino acid ***i*** observed for a codon ***j***. In contrast to *S. cerevisiae*, we find the Tryptophan codon TGG misincorporated as a variety of amino acids. **f)** Codon specific error detection rate in *E. coli*. We observe similar variability of error rates between synonymous codons and datasets to *S. cerevisiae*. For most amino acids, *E. coli* and *S. cerevisiae* share the codon with the highest fidelity. Notable exceptions are Leucine and Valine (thick black line indicates median, box indicates 25$^{th}$ and 75$^{th}$ percentiles, whiskers indicate 1.5 times the inter-quartile range).

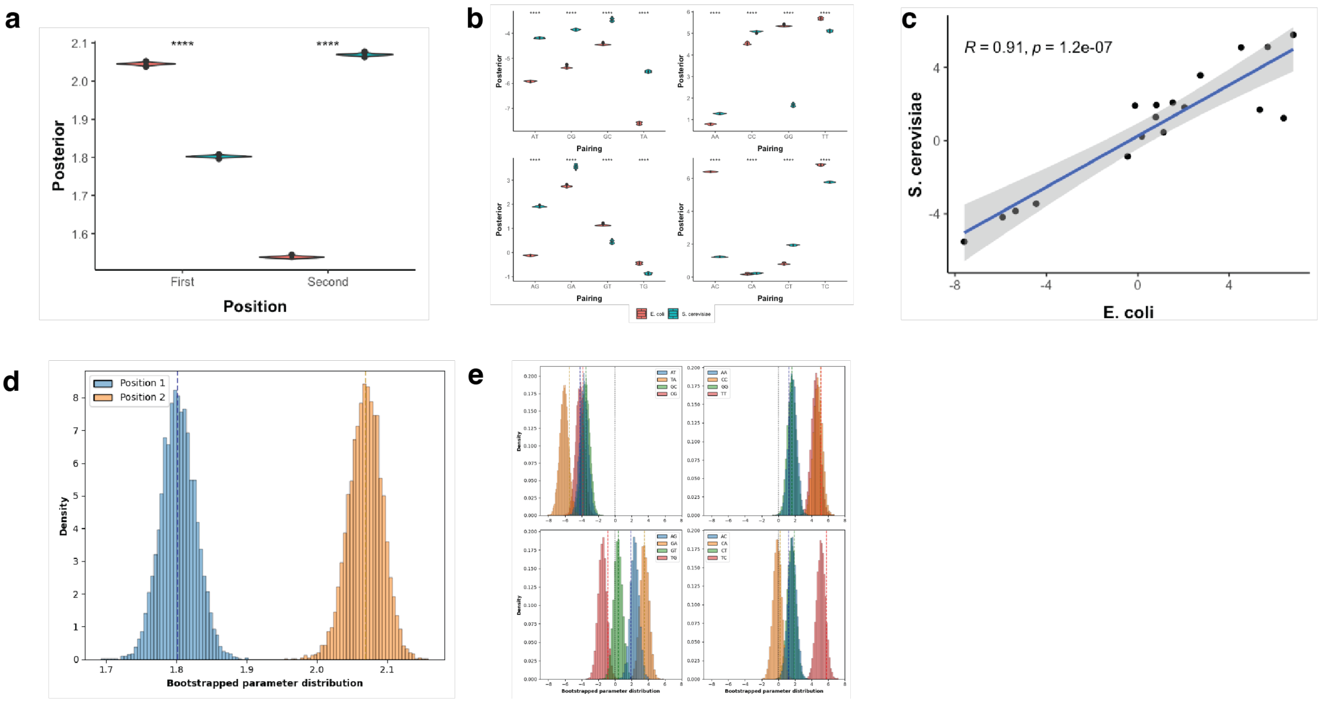


**Figure S2: Parameter bootstrap and model parameter posterior distributions between *S. cerevisiae* and *E. coli*** **a)** Distribution of position posterior means. **b)** Distribution of wobble parameter posterior means. **a, b)** Dashed line indicates model fit. **c)** Distribution of the position posteriors. **d)** Distribution of wobble parameter posteriors. **c, d)** All distributions show a significant difference in their mean values between *S. cerevisiae* and *E. coli* (Kruskal-Wallis test, $\rho < 0.0001, n = 500$).

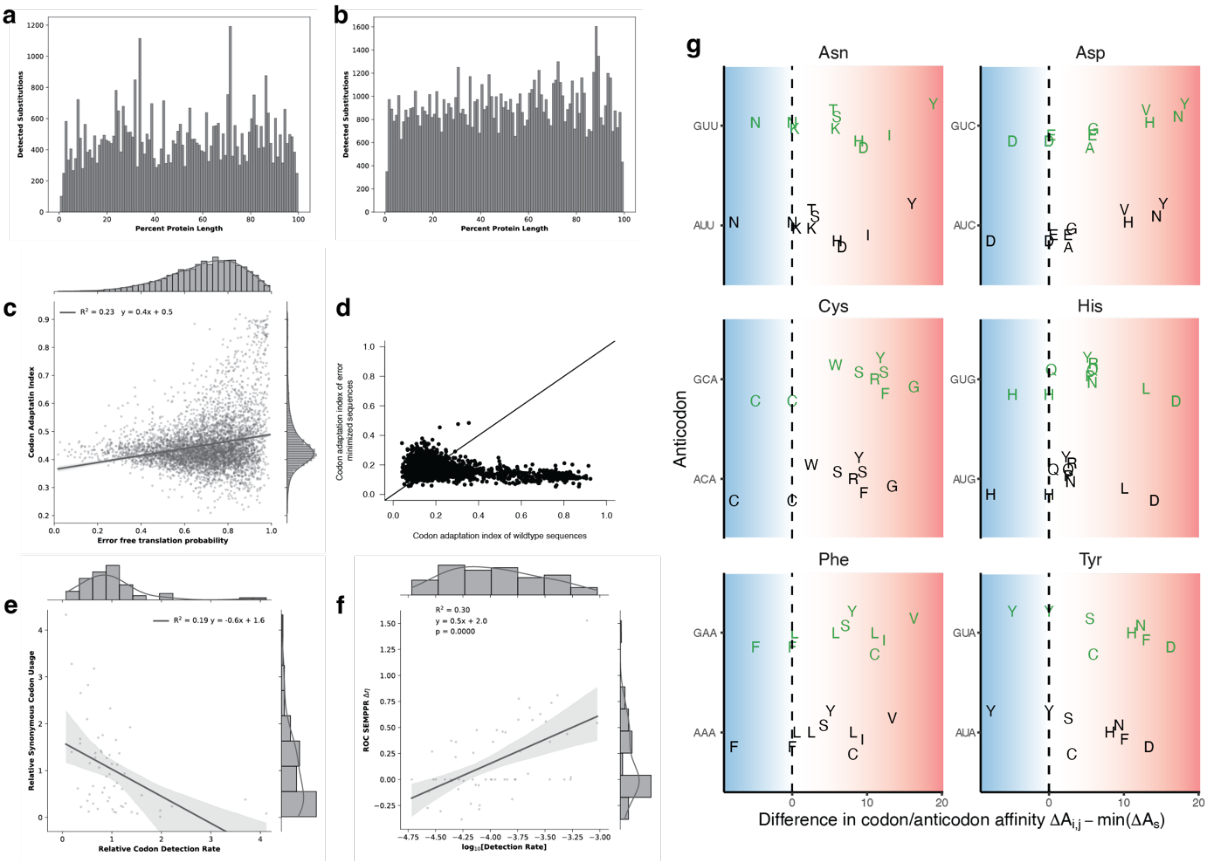


**Figure S3*: Exploring effects of codon/anticodon usage on translation efficiency fidelity.*** **a)** *S. cerevisiae* shows a mostly uniform distribution of amino acid misincorporations across all proteins in the proteome. **b)** *E. coli* also shows a mostly uniform distribution. In contrast to *S. cerevisiae*, however, a noticeable drop-off in detected misincorporations can be observed in the last bin; potentially explained by the lack of tryptic peptides at the end of a protein. **a, b)** Each bin represents one percent protein length. **c)** Codon adaptation index (CAI) is only weakly correlated with error free translation probability. While genes with high CAI are more likely to be translated error free, the opposite does not hold as many proteins with low CAI show high fidelity. **d)** Comparing the codon adaptation index (CAI) wild-type and fidelity optimized codon sequences reveals a drop in sequence adaptation for sequences with a high CAI. Wild-type sequences with a low CAI and a codon usage likely dominated by mutation tend to improve their codon adaptation. **e)** Relative synonymous codon usage describing the preferred codon usage of the most highly expressed genes (top 5 %) is negatively correlated with the relative codon detection rate indicating that frequent codons are translated more accurately. **f)** ROC-SEMPPR's $\Delta\eta$ values, describing a codons translation inefficiency (increasing $\Delta\eta$ indicates inefficient codons) are positively correlated with a codons error detection rate. Indicating that inefficiently translated codons tend to have a higher error rate. **g)** Impact of the tRNA pool on translational errors. Difference in codon/anticodon affinity for interactions with up to one codon/anticodon mismatch, for all two-codon amino acids with only one tRNA present in *E. coli*. The ability of the ribosome to discriminate between correct and incorrect binding events is diminished if we assume that the missing tRNA (black) would be present rather than the tRNA naturally present (green).

| Parameter | Posterior Mean | Bootstrap Mean | Empirical P-value |
|---|---|---|---|
| Position 1 | 1.802 | 1.719 | 0.505 |
| Position 2 | 2.069 | 2.069 | 0.509 |
| AA | 1.278 | 1.719 | 0.802 |
| AC | 1.225 | 1.673 | 0.805 |
| AG | 1.902 | 2.352 | 0.806 |
| CA | 0.214 | -0.121 | 0.261 |
| CC | 5.068 | 4.734 | 0.268 |
| CT | 1.919 | 1.586 | 0.266 |
| TC | 5.779 | 5.136 | 0.115 |
| TT | 5.124 | 4.486 | 0.109 |
| TG | -0.858 | -1.484 | 0.113 |
| GA | 3.535 | 3.519 | 0.483 |
| GT | 0.412 | 0.388 | 0.478 |
| GG | 1.645 | 1.611 | 0.473 |
| AT | -4.186 | -3.737 | 0.808 |
| TA | -5.509 | -6.109 | 0.133 |
| GC | -3.487 | -3.513 | 0.479 |
| CG | -3.866 | -4.201 | 0.260 |

**Table S1: Model parameter posterior means with empirical p-value based on 10.000 bootstrap samples for S. cerevisiae.** Empirical p-value was calculates as $(r+1)/(n+1)$ were $r$ is the number of sample greater than the posterior mean, and $n$ is the total number of samples[68].
